# Integrated Molecular Profiling Studies to Characterize the Cellular Origins of High-Grade Serous Ovarian Cancer

**DOI:** 10.1101/330597

**Authors:** Kate Lawrenson, Marcos A.S. Fonseca, Felipe Segato, Janet M. Lee, Rosario I. Corona, Ji-Heui Seo, Simon Coetzee, Yvonne G. Lin, Tanja Pejovic, Paulette Mhawech-Fauceglia, Ronny Drapkin, Beth Y. Karlan, Dennis J. Hazelett, Matthew L. Freedman, Simon A. Gayther, Houtan Noushmehr

## Abstract

Historically, high-grade serous ovarian cancers (HGSOCs) were thought to arise from ovarian surface epithelial cells (OSECs) but recent data implicate fallopian tube secretory epithelial cells (FTSECs) as the major precursor. We performed transcriptomic and epigenomic profiling to characterize molecular similarities between OSECs, FTSECs and HGSOCs. Transcriptomic signatures of FTSECs were preserved in most HGSOCs reinforcing FTSECs as the predominant cell-of-origin; though an OSEC-like signature was associated with increased chemosensitivity (*P*_*adj*_ = 0.03) and was enriched in proliferative-type tumors, suggesting a dualistic model for HGSOC origins. More super-enhancers (SEs) were shared between FTSECs and HGSOCs than between OSECS and HGSOCs (*P* < 2.2 × 10^−16^). SOX18, ELF3 and EHF transcription factors (TFs) coincided with HGSOC SEs and represent putative novel drivers of tumor development. Our integrative analyses support a predominantly fallopian origin for HGSOCs and indicate tumorigenesis may be driven by different TFs according to cell-of-origin.

---

Invasive epithelial ovarian cancers are a heterogenous group of tumors comprising several major histological subtypes: high-grade serous, low-grade serous, endometrioid, clear cell and mucinous. High-grade serous ovarian cancer (HGSOC) is the most common subtype, comprising around two-thirds of all invasive cases. Our understanding of the cellular origins of HGSOC and key transcription factor networks deregulated during HGSOC development has been restricted by the lack of substantial molecular profiling data for the suggested precursor tissues, ovarian surface epithelial cells (OSECs) and fallopian tube secretory epithelial cells (FTSECs). Historically, HGSOCs were thought to arise from OSECs, an atypical epithelial cell type with mesothelial features and inherent phenotypic plasticity and heterogeneity ^1,2^.However, examples of early-stage ovarian carcinoma arising from OSECs *in vivo* are scare, and the discovery of occult carcinomas in the fallopian tubes of *BRCA1/2* mutation carriers provided evidence to support an alternative hypothesis that the fallopian epithelium harbors the cell-of-origin for HGSOC ^3–7^. Subsequent studies support a tubal origin for HGSOCs not associated with highly penetrant gene mutations, suggesting that HGSOCs arise from FTSECs in at least 50% of cases ^8–10^. However, in a proportion of cases there remains no evidence of fallopian involvement, which raises the possibility that HGSOCs could originate from more than one tissue type. The large-scale molecular data sets required to fully evaluate the relationships of OSEC and FTSECs to HGSOC have been lacking, and so in the current study we established transcriptome-wide signatures in 188 normal cells representing these two putative origins of HGSOC. We used machine learning to quantify the similarities between OSECs and FTSECs to HGSOCs transcriptomes as a molecular approach to characterizing disease origins. In addition, we integrated transcriptomic data with epigenetic maps for these cell types, to characterize tissue-specific super-enhancer landscapes, and to identify putative transcription factors driving transcriptional deregulation during serous ovarian tumorigenesis.

## Results

### Expression profiling of putative ovarian cancer precursor cells

One approach to exploring cellular origins of cancer is to quantify similarities and differences between molecular signatures of tumors and the proposed tissues of origin ^11^; based on the hypothesis that the molecular blueprint of the normal precursor cell is maintained in the developed tumor. We applied this approach to explore HGSOC origin by performing RNA-sequencing (RNA-seq) on an extensive series of normal cells isolated from the two major proposed precursor tissue types, ovarian surface epithelial cells (OSECs; n = 114) and fallopian tube secretory epithelial cells (FTSECs; n = 74). Cell samples were derived from 132 individuals, with both OSECs and FTSECs isolated from the same patient in 56 cases ^12,13^ (Supplementary Table 1). For one sample, we replicated the RNA sequencing for quality control purposes; expression profiles from this samples were highly correlated (Pearson’s correlation r = 0.98) (Supplementary Fig. 1). We found no associations with experimental or epidemiological variables (where available), including sample preparation patient age or ethnicity (data not shown).

**Figure 1.**
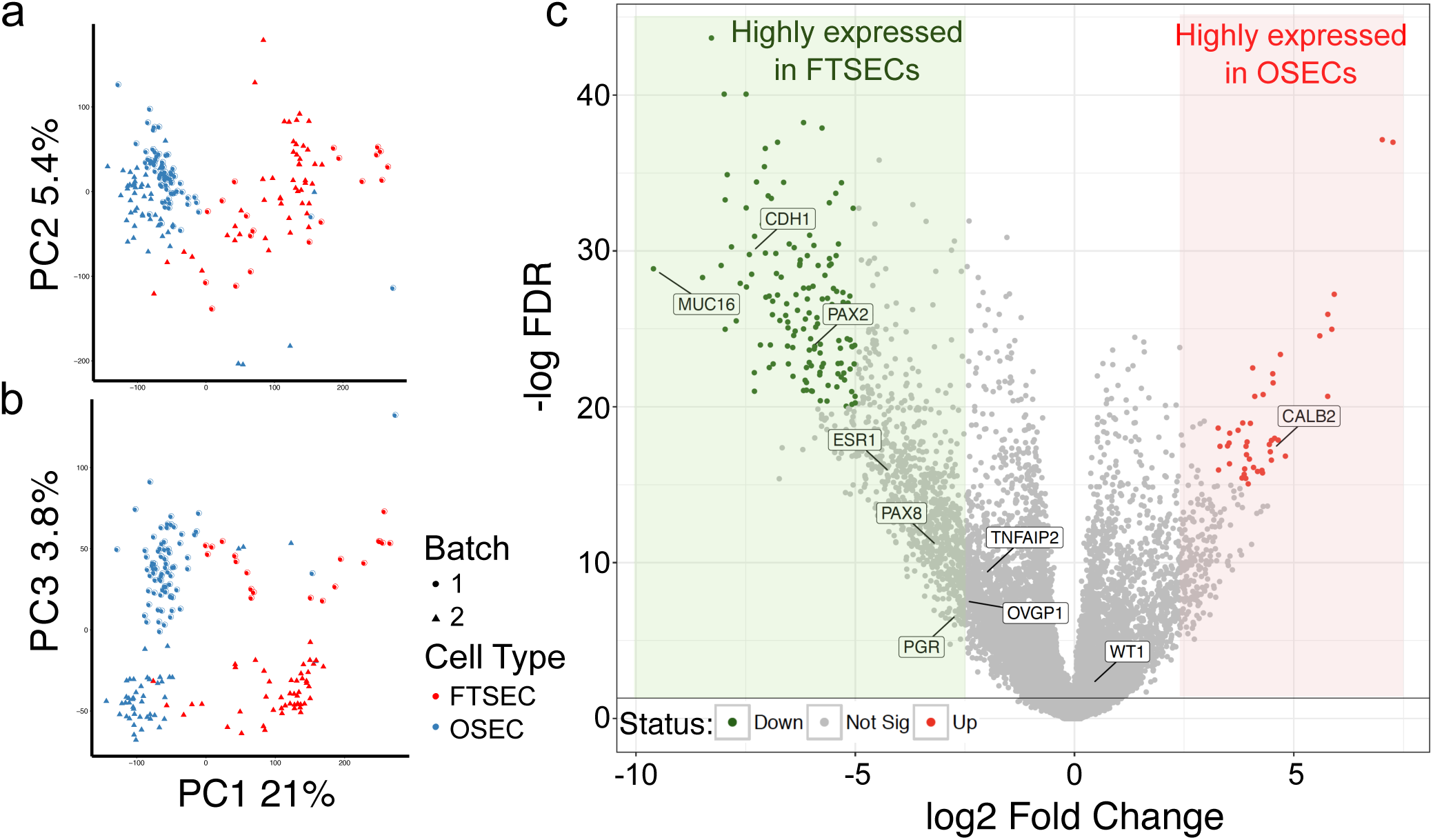
Transcriptomic profiling of OSECs and FTSECs. (a & b) Principal component analysis (PCA) of RNA-seq profiles of OSECs and FTSECs. OSEC samples tend to cluster more tightly together whereas FTSEC samples show more diffuse clustering. This suggests greater inter-patient heterogeneity between for FTSEC samples. (c) Volcano plot illustrating differential gene expression between OSEC and FTSEC samples. Known cell-type specific markers for each cell type are indicated.

Using transcriptome-wide principal component analysis (PCA) to compare expression signatures between OSECs and FTSECs, the two cell types largely stratified according to their molecular profiles (Fig. 1a,b). We identified the 87 genes that were the most differentially expressed between OSECs and FTSECs (absolute log_2_ fold change [FC] > 2, P_adj_ = 10^−30^, Fig. 1d, Supplementary Table 2). These included *MUC16* (which encodes ovarian cancer screening marker CA125) and *CDH1* (E-Cadherin), genes known to be differentially expressed between these cell types (Fig. 1c, Supplementary Fig. 2). Novel, highly overexpressed differentially expressed genes (DEGs) in OSECs compared to FTSECs included *GATA4* (FC = 7.1, P_adj_ = 3.78×10^−42^) and *NR5A1* (FC = 6.7, P_adj_ = 2.59×10^−39^); both transcriptional activators potentially involved in differentiation of the OSEC phenotype. Novel DEGs highly expressed in FTSECs compared to OSECs include genes that encode the cell surface or secreted proteins *MMP7* (FC = 9.9, P_adj_ = 1.87×10^−31^), *CLIC5* (FC = 8.36, P_adj_ = 5.14×10^−49^), *TACSTD2* (FC = 8.21, P_adj_ = 1.2×10^−42^) and *CFTR* (FC = 8.15, P_adj_ = 7.35×10^−31^).

**Figure 2.**
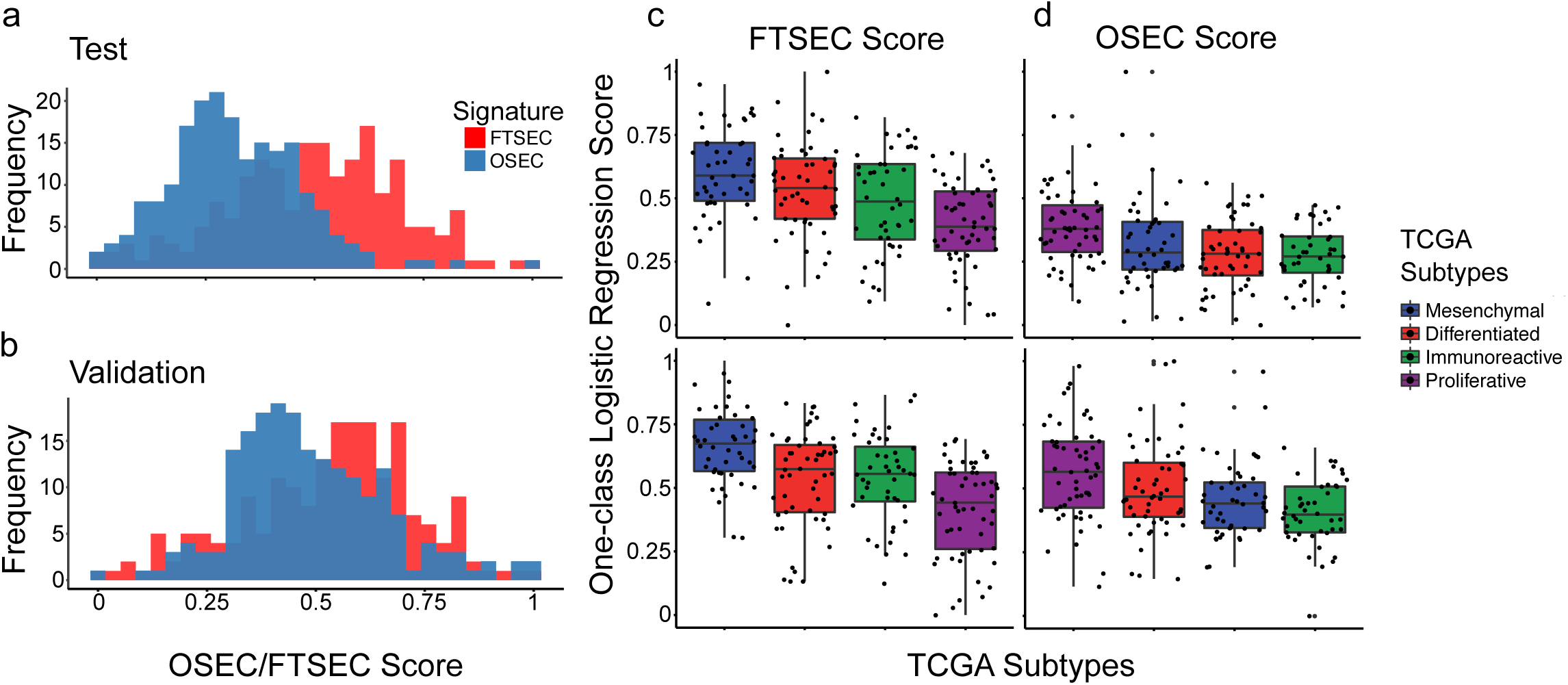
One-class logistic regression (OCLR) predictors of the cellular origins of HGSOC. (a & b) OSEC and FTSEC whole transcriptomic signatures were developed and compared to whole transcriptomic signatures of 394 primary HGSOCs publicly available from TCGA project. TCGA analyses were divided into (a) test (n=197) and (b) validation (n=197) sets of tumors. In both test and validation sets, FTSEC score tended to be higher in HGSOCs than OSEC scores. The dashed line indicates a score of 0.5. (c & d) FTSEC and OSEC signatures were compared across HGSOC molecular subgroups. FTSEC score was highest in the mesenchymal subgroup in both test and validation sets; (c) OSECs were highest in the proliferative subgroup.

### Transcriptomic Relationships between OSECs, FTSECs and HGSOCs

We applied machine learning algorithms to evaluate the molecular relationships between transcriptomic profiles of OSECs and FTSECs and 394 primary HGSOCs profiled by The Cancer Genome Atlas (TCGA). To correct for differences in read depth and RNA-seq methods between this study and TCGA, we aligned, batch corrected and normalized all three data sets (OSEC, FTSEC and TCGA) together (see Methods). We first defined cell-type specific signatures of OSECs and FTSECs and then applied a One-class logistic regression (OCLR) methodology ^14^. Area under the curve (AUC) statistics generated using a leave-one-out approach indicated that the OCLR models performed with high specificity (average AUC for OSECs = 0.99 and for FTSECs = 0.97). OCLR models provide a score for each sample and for each category, which is rescaled between zero and one, where zero implies no similarity and one implies high similarity. We applied the OCLR models to HGSOCs to evaluate which cell type - OSECs or FTSECs - was most highly correlated with HGSOC, thereby indicating which cell type is the most likely cell-of-origin. HGSOC samples were randomized and divided into two equally sized groups (n = 197), designated the training set and the validation set. Each set included similar numbers of the four HGSOC molecular subgroups - differentiated, immunoreactive, mesenchymal and proliferative, classified based on gene expression signatures. In both the training and validation data sets, we observed a greater proportion of HGSOCs with higher FTSEC scores compared to OSEC scores. In the training set 103/197 tumors (52%) had an FTSEC score > 0.5, while only 20/197 tumors (10%) had an OSEC score > 0.5. In the validation set 124/197 tumors (63%) and 82/197 tumors (42%) had FTSEC and OSEC scores > 0.5, respectively (Fig.2a,b). Taken together, these data indicate that the molecular signatures of HGSOCs are more similar to FTSECs than OSECs. There was a weak negative correlation between tumor FTSEC and OSEC scores (Supplementary Fig. 3) (Pearson’s product-moment correlation = -0.16, *P* = 0.002). Consistent with our machine learning observations, in a PCA performed using all expressed genes, FTSECs cluster more closely to HGSOCs than OSECs (Supplementary Fig. 4).

**Figure 3.**
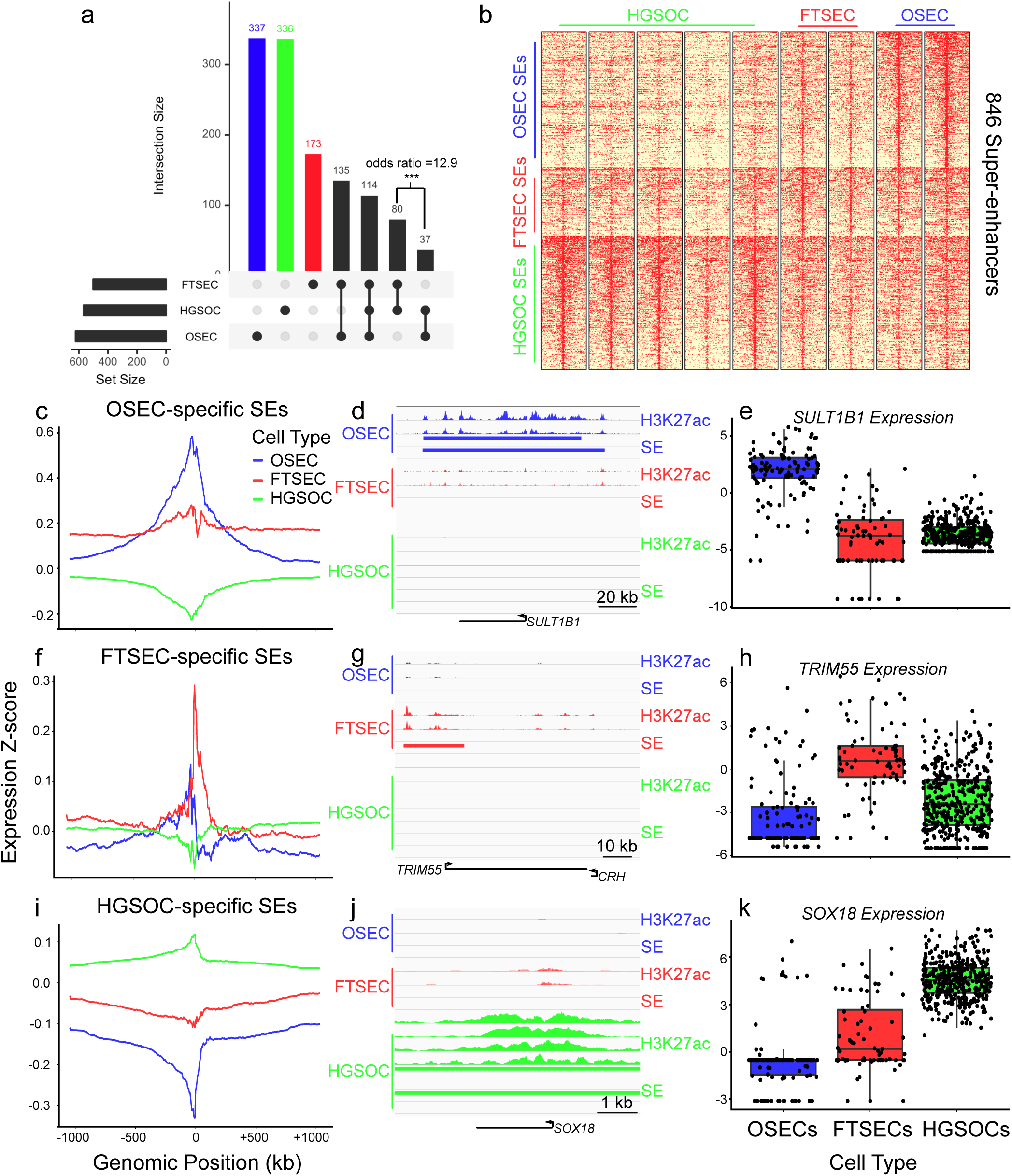
Super-enhancer-gene relationships in OSECs, FTSECs and HGSOCs. (a) Pseudo Venn diagram of the SE catalogue of OSECs, FTSECs and HGSOCs, showing SE intersections across the three tissue types. Although fewer SEs were catalogued in FTSECs compared to OSECs and HGSOCs, significantly more SEs are shared between FTSECs and HGSOCs than between OSECs and HGSOCs (Fishers Exact Test, p < 2.2 × 10^−16^). (b) The landscape of cell-type specific SEs across OSECs, FTSECs and HGSOCs. For OSECs and FTSECs, H3K27ac ChIP-seq data generated for two independent immortalized normal lines per cell type were used to identify SEs. For HGSOCs, H3K27ac ChIP-seq data were generated for five different primary HGSOCs. (c-k) Tissue-specific SEs associated with elevated gene expression in cis in a cell-type specific manner. (c-e), OSEC-specific SEs, (f-h) FTSEC-specific SEs, (i-k) HGSOC-specific SEs. (c,f,i) H3K27ac ChIPseq data were integrated with RNA-seq data for each tissue type. Average gene expression in 114 OSECs, 74 FTSECs and 394 HGSOCs is shown for regions centered on cell-type specific SEs; (d,g,j) Representative loci displaying tissue-specific SE deposition for each tissue type; (e,h,k) Box plots illustrating differential gene expression between tissue types for candidate, cell type specific cis-regulated genes. The associated gene consistently displays higher expression in the SE-positive tissue type.

**Figure 4.**
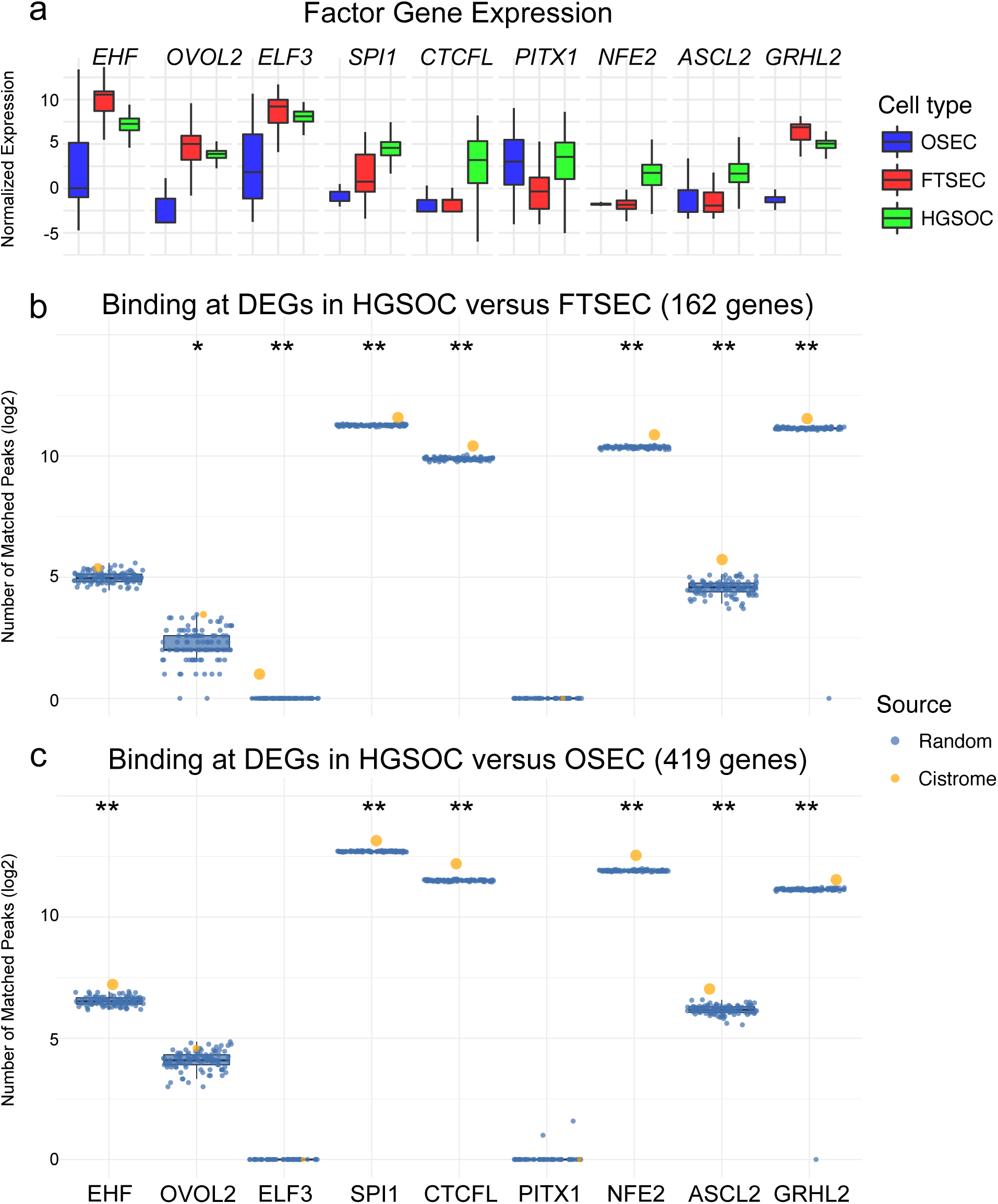
Novel transcriptional regulators implicated in HGSOC development. (a) Transcriptional regulators highly expressed in HGSOC. The number of differentially expressed genes in HGSOCs compared to (b) FTSECs and (c) OSECs that localize with ChIP-seq peaks for each factor (orange) and factor-specific matched random peaks (blue). * *P* > 0.05, ** *P* > 0.01.

We also evaluated if FTSEC and OSEC scores correlated with clinical and molecular features of HGSOCs ^15,16^. In both the training and validation data sets, mesenchymal-type HGSOCs had significantly higher FTSEC OCLR scores (*P*_*adj*_ < 0.02 in the training and validation cohort, Fig. 2c, *P*_*adj*_ = 8×10^−4^ in a meta-analysis of all 394 HGSOCs); this subgroup of HGSOCs exhibit the shortest survival times ^15^. By contrast, proliferative type HGSOCs had significantly larger OSEC scores (*P*_*adj*_ < 0.001 in the training and validation cohort, *P*_*adj*_ = 2×10^−4^ in a meta-analysis) (Fig. 2d) suggesting that OSECs may be the more likely cell-of origin for this subgroup of HGSOCs. Taken together, these data are consistent with the hypothesis that HGSOCs derive from more than one tissue type ^17–19^. Finally, we tested for associations between FTSEC and OSEC OCLR scores and patient age, tumor stage, tumor grade, chemoresponse and debulking status (Supplementary Figs. 5,6). We found no significant associations for FTSECs, but tumors with high OSEC scores were associated with older age at diagnosis (*P*_*adj*_ = 0.005, normalized enrichment score = 1.6). Higher OSEC score was also associated with increased sensitivity to chemotherapy (*P*_*adj*_ = 0.03, normalized enrichment score = 1.5) (Supplementary Fig. 6).

### Identifying Epigenomic and Transcriptomic Interactions

We used chromatin immunoprecipitation sequencing (ChIP-seq) to characterize epigenomic landscapes in OSECs, FTSECs and HGSOCs, focusing on super-enhancers (SEs), which are dense clusters of highly active chromatin that typically localize with master regulators of cellular identity. We generated ChIP-seq data for acetylated H3K27ac in five primary HGSOC tissues; H3K27ac ChIP-seq data for normal OSEC and FTSEC cells have been reported previously ^20^. H3K27ac profiles were used to map SEs in each tissue type. OSECs and HGSOCs had the largest numbers of cell-type specific SEs (n=337 and n=336, respectively). FTSECs had the lowest number of unique SEs (n=173); this was due largely to the fact that there were significantly more SEs shared between FTSECs and HGSOCs (n=80) than between OSECS and HGSOCs (n=37) (odds ratio = 12.9, Fishers Exact Test, p < 2.2 × 10^−16^, Fig. 3a,b). Using the transcriptomic data described above, we verified tissue-specific overexpression of genes located at tissue-relevant SE loci (Fig. 3c,f,i), and identified novel candidate target genes regulated by SEs in each cell type. These included *SULT1B1* in OSECs (Fig. 3d,e); the Tripartite Motif Containing 55 (*TRIM55*) gene in FTSECs (Fig. 3g,h) and the *SOX18* transcription factor in HGSOCs (Fig. 3j,k). Of particular note, the *PAX8* transcription factor was overexpressed in both FTSECs and HGSOCs and coincides with a SE detected in both cell types at this locus; PAX8 is a well-established biomarker that is ubiquitously expressed in FTSECs and is overexpressed in the majority of primary HGSOCs ^21–23^ (Supplementary Fig. 7).

Finally, we performed a targeted analysis of genes associated with DNA binding, transcription factor activity and chromatin remodeling to identify putative novel drivers of transcriptional deregulation during HGSOC development from FTSECs and/or OSECs. For nine of the most overexpressed transcriptional regulators in HGSOCs (Fig. 4a), high-quality ChIP-seq data were available from cistrome.org. Using these ChIP-seq data sets we quantified how many of the most differentially expressed genes in HGSOCs were located near to (within 50 kbp) a factor-specific peak compared to matched random peaks (see Methods). Factor-specific peaks for regulators including SPI1, CTCFL, NFE2, ASCL2 and GRHL2 were more numerous in the vicinity of HGSOC DEGs (*P* < 10^−30^ for a comparison to FTSECs and *P* < 10^−50^ for a comparison to OSECs) than randomly generated matched sets of background peaks (100 iterations, *P* <0.01 Fig. 4b,c; genes located close to factor peaks are listed in Supplementary Tables 3,4). Binding sites for ELF3, a factor highly expressed in both FTSECs and HGSOCs but lowly expressed in OSECs were specifically enriched near to genes differentially expressed between FTSECs and HGSOCs (*P* = 0.009, Fig. 4b), with no evidence of enrichment in the set of genes differentially expressed in the development of HGSOC from OSECs (*P* = 1, Fig. 4c). Conversely EHF binding sites were associated with DEGs in a comparison of HGSOCs to OSECs but not FTSECs (*P* = 0.009 and *P* = 0.06, respectively). Notably, both ELF3 and EHF reside at SEs in HGSOC (Supplementary Figs. 8,9). Collectively these factors represent novel drivers of transcriptional reprogramming in HGSOC, with ELF3 likely to be specific to the transformation of FTSECs, and EHF a putative driver of HGSOC development from OSECs.

## Discussion

There remains debate about the cellular origins of high-grade serous ovarian cancer. Over the last few years the fallopian tube, and specifically the secretory epithelial cell component (FTSECs), has emerged as a likely origin. This is based on the identification of early stage precursor lesions in the fallopian tube that express secretory cell lineage markers and harbor TP53 mutations, particularly in fallopian tube fimbriae ^3,4,8,10,24^. In addition, *in vitro* and *in vivo* models studies ^12,25,26^ support FTSECs as a major cell-of-origin for HGSOC and salpingectomy (surgical removal of fallopian tubes but not the ovaries) can reduce the risk of ovarian cancer by around 35% or more ^27^. However, the current data do not implicate FTSECs as precursors of every HGSOC, in particular in non-familial cases (i.e. non-carriers of *BRCA1/BRCA2* mutations). This suggests that alternative cellular precursors may exist, which supported the present re-evaluation of ovarian surface epithelial cells (OSECs), historically thought to the precursor of HGSOCs.Collectively, in a cohort of almost 400 HGSOCs, we found greater similarities in transcriptomic signatures between FTSECs and HGSOCs than between OSECs and HGSOCs. Primary HGSOCs can be sub-stratified into 4 different molecular groups based on mRNA expression profiles and high FTSEC scores were enriched in mesenchymal-type HGSOCs, consistent with previous observations indicating that fallopian-like HGSOCs are associated with poorer outcomes ^28^. OSEC profiles were most closely related to the proliferative-type HGSOCs associated with better outcomes. These data imply that HGSOC molecular subgroups may have different precursors, with OSECs the putative cell-of-origin for the proliferative subgroup. While our results add to a growing body of evidence supporting FTSECs as the predominant cell-of-origin for HGSOC ^9,19,29^, there remains a substantial and often overlooked body of evidence that supports OSECs as alternative precursors in a subgroup of cases. OSECs can express many HGSOC markers, including PAX8 ^2,30^. OSECs from primary ovaries and OSEC cultures from women at high-risk of ovarian cancer show phenotypic differences compared to OSEC cultures from non-inherited cases, with OSECs from high-risk women more committed to an epithelial phenotype and maintaining expression of CA125 longer than OSECs from non-high-risk women ^31^. In addition, occult cancers have been detected in the ovaries of women undergoing prophylactic risk reducing oophorectomy and can occur without evidence of lesions in the fallopian tube ^32^. Our results are also consistent with a recent detailed analysis of almost 60 ovaries which documented evidence for metaplasia of ovarian epithelium to a Müllerian phenotype, and proposed that this could be an early step in the development of serous tumors ^2^.

While our study shows clear differences between the transcriptomic profiles of OSECs and FTSECs, there are some caveats to the study design that could influence interpretation of our findings. One of these is the potential impact of short term culture on the gene expression profiles of primary OSECs and FTSECs. Neither cell type is readily amenable to microdissection from primary tissues - OSECs are a scarce population on the surface of the ovary and FTSECs are intercalated with ciliated cells not currently thought to be precursors of serous carcinogenesis; thus culturing cells was our preferred approach to obtaining sufficient purified OSEC/FTSEC cell material on which to perform in-depth molecular profiling studies without having to pool samples across patients. To limit artifacts induced by cell culture, cells were not genetically modified and the procedures for primary culture were optimized for each cell type specifically.

Moreover, our results are closely reflected by results of a recent molecular profiling study which compared an independent cohort of HGSOCs to signatures derived from a small cohort of fallopian (n=3 pooled samples) or ovarian brushing (n=4 pooled samples) and observed similar trends: that the majority of HGSOCs correlate with fallopian signatures but a subset of cases show higher correlation with OSEC signatures ^29^; suggesting that short-term culture did not substantially influence our analyses.

Little is known about the key transcription factors (TFs) driving oncogenesis in HGSOC. PAX8 is a TF known to be highly expressed in FTSECs and functionally involved in disease development ^22,30,33–35^. We found that PAX8 is marked by a super-enhancer in both FTSECs and HGSOCs. We also identified ELF3 and EHF as putative drivers of the transformation of FTSECs and OSECs respectively, both of which have been implicated in ovarian cancer development ^36,37^. Of the four genes highly expressed in OSECs, two were transcriptional regulators: *GATA4* is a developmental zinc finger TF and *NR5A1* is a transcriptional activator involved in sex determination and differentiation of steroidogenic tissues. Both *GATA4* and *NR5A1* were highly expressed in proliferative type HGSOCs, suggesting these TF networks may specifically be involved in the development of this tumor subgroup.

Compared to previous studies, these investigations represent a significant advance in both scale and scope of the molecular profiling performed ^28^. While this study focused on the HGSOC subtype of ovarian cancer, where there is good evidence for the likely precursor cell types, a similar approach could also be used to investigate the cells of origin of other histotypes; OSECs, FTSECs and other Müllerian epithelia should also be comprehensively evaluated as putative cells-of-origin for low-grade serous, mucinous, clear cell and endometrioid ovarian cancers. In summary, this study provides novel insights into the etiology and biology of likely precursors of HGSOC, ovarian surface and fallopian tube secretory epithelial cells. In addition we have described for the first time, the putative transcriptional regulators that are associated with these cell types and their role in HGSOC development; these represent candidate clinical biomarkers and novel therapeutic targets for this disease.

## Methods

### Patient consent and IRB approval

All specimens were collected from University College Hospital (London, UK), LAC + USC Medical Center (Los Angeles, CA, USA) and Oregon Health & Science University (Portland, OR, USA). All were collected with informed patient consent and Institutional Review Board approval.

### Sample collection, RNA extraction and RNA sequencing

OSECs and FTSECs were harvested from women diagnosed with ovarian, uterine or cervical cancer, and were histologically normal. Short-term cultures were established as previously described^12,13^. Briefly, OSECs were harvested using a cytobrush and cultured in NOSE-CM media containing 15% fetal bovine serum (FBS, Hyclone), 34 µg ml^−1^ bovine pituitary extract, 10 ng ml^−1^ epidermal growth factor (Life Technologies), 5 µg ml^−1^ insulin and 500 ng ml^−1^ hydrocortisone (Sigma-Aldrich). FTSECs were harvested by Pronase/DNase I digestion (Roche and Sigma-Aldrich, respectively) for 48-72 hours at 4°C and cultured on collagen I (Sigma-Aldrich) using DMEM/F12 base media supplemented with 2% Ultroser G (Pall Corporation). At ∼80% confluency, cells were lysed using the QIAzol reagent and RNA extracted using the RNeasy Mini kit (both QIAgen). RNA sequencing was performed by the University of Southern California Epigenome Core Facility.

### RNAseq data processing and QC

All data analysis was performed using ‘R’ and ‘Bioconductor’, and packages therein. RNAseq data for 394 HGSOC samples was obtained from The Cancer Genome Atlas (TCGA) data portal as protected data (raw sequencing, fastq files) and downloaded via CGHub’s geneTorrent. Data was aligned to a reference genome (hg19) using STAR. And quality control of aligned samples performed using RSeQC. GC bias and batch effect corrections were performed using EDASeq and ‘sva’. To adjust for batch effects we used an empirical Bayes framework (comBat), available in ‘sva’.

### Differential gene expression analyses

After normalization, the data matrix contained 21,071 genes. Parametric statistics (Student’s T-test) and supervised hierarchical clustering were performed to identify genes differentially expressed in pairwise comparisons of two groups of interest (OSEC, FTSEC and HGSOC). P-values were adjusted using Benjamini–Hochberg step-up procedure.

### Machine learning analyses

We applied a machine learning approach to define a probabilistic score associated to both normal cell types, and infer tumor origins. A One-class classifier was selected as this method can handle non-traditional supervised scenarios where no negative class can be defined. The classifiers were implemented by the gelnet R-package version 1.2.1. Data were mean centered considering all samples together, then each cell type used separately to train and test the models. To train the OSEC model we considered all OSEC samples, with a coefficient for the L1-norm penalty equal to 0 and coefficient for the L2-norm penalty equal to 1 as arguments of gelnet function. We then evaluated the model performance through leave-one-out procedure where the left-out OSEC sample was mixed into FTSEC sample background. The accuracy was evaluated via the Area Under the ROC curve method, with 99% of OSEC samples correctly predicted, on average, and 97% of FTSECs correctly predicted. We then tested the model prediction behaviour when applied to HGSOCs from TCGA. We took advantage of the fast gene set enrichment analysis (fgsea, version 1.2.1, http://bioconductor.org/packages/fgsea/) method to evaluate enrichment of clinical attributes across the tumor OCLR scores from both FTSEC and OSEC models. We applied the fgsea function with the parameter nperm equal to 10,000.

### Tissue ChIP-seq in HGSOC specimens

Tissue ChIP-seq was performed based on the methods described in Pomerantz *et al* 2015.^38^. One 3 mm core was isolated from an epithelial-rich portion of tumor, and pulverized using the Covaris CryoPrep system (Covaris, Woburn, MA), set to an intensity of 4. Tissues were fixed using 1% formaldehyde (Thermo fisher, Waltham, MA) for 10 minutes at room temperature. Fixation was quenched with 125 mM glycine and samples were rinsed with cold PBS before a 10 minute lysis in a buffer containing 50 mM Tris, 10mM EDTA, 1% SDS with protease inhibitor). Chromatin was sheared to 300–500 base pairs and 5 volumes dilution buffer (1% Triton X-100, 2 mM EDTA, 150 mM NaCl, 20 mM Tris HCl pH 8.1) added. Each sample was incubated with 1 µg H3K27ac antibody (DiAGenode, C15410196, Denville, NJ) coupled with protein A and protein G beads (Life Technologies, Carlsbad, CA) at 4°C overnight. Immunoprecipitated chromatin was washed with RIPA buffer (0.05M HEPES pH 7.6, 1 mM EDTA, 0.7% Na Deoxycholate, 1% NP-40, 0.5M LiCl) five times and rinsed with TE buffer (pH 8.0) once. The sample was resuspended in elution buffer (50mM Tris, 10mM EDTA, 1% SDS), treated with RNase for 30 minutes at 37°C, and incubated with proteinase K overnight at 65°C. Sample DNA and 1% input were extracted, and sequencing libraries prepared using the ThruPLEX-FD Prep Kit (Rubicon Genomics, Ann Arbor, MI). Libraries were sequenced using 75-base pair single reads on the Illumina platform (Illumina, San Diego, CA) at the Dana-Farber Cancer Institute.

### ChIP-seq data analysis

The AQUAS pipeline (https://github.com/kundajelab/chipseq_pipeline) was used to processed ChIP-seq data. Reads were aligned to the reference human genome (hg19), filtered by read quality and duplicate reads removed. macs2 (https://pypi.python.org/pypi/MACS2) was used for peak calling. For the cell lines, two technical replicates were generated and the final peaks were obtained using a naive overlap approach, where the peaks are included if they overlap more than 50% between the two technical replicates. We have previously described H3K27ac ChIP-seq for immortalized OSEC and FTSEC lines ^20^. Immortalized OSECs have been previously show to be representative of unmodified cells ^39^. We verified that the expression profiled of immortalized FTSECs used in this study clustered with primary FTSECs (Supplementary Figure 10). After alignment, homer (http://homer.ucsd.edu/homer/) was used to identify super-enhancers, using a super slope parameter of 2 and a minimum distance of ten thousand. For defining a set of HGSOC SEs, we selected SEs that were called in at least two HGSOC samples. For the FTSEC set of SEs, SEs were called individually in each technical replicate, then, all the SEs that overlapped both technical replicates within the same cell line (FTSEC33 or FTSEC246) were selected to get the union set. We used a similar approach to get the union set of SEs for the OSEC cell lines.

### Cistrome TF analyses

From the Cistrome database (cistrome.org), we selected and downloaded BED peak files for experiments passing 3/6 quality control filters. Data sets used are listed in the Online Methods. The following data sets were included: HOXC9: GSM848788, GSM848789; LXH2: GSM1208644,GSM1567049; GLIS1: ENCSR482BBZ, GSM1208752, GSM2026832, GSM803384; GATA4: ENCSR590CNM, GSM1505646, GSM1505647, GSM1505651, GSM1505657; HNF1B: GSM1239408, GSM1505680, GSM1505681, GSM1505682, GSM1505683; EHF: GSM1208609, GSM1548071, GSM1548072; OVOL2: GSM1239518; ELF3: GSM1208732; SPI1: GSM1010843, GSM1370280, GSM1370292, GSM1421017, GSM1642769, GSM1703900; CTCFL: GSM1239559, GSM1817655, GSM1817659, GSM1817663, GSM1817668, GSM1817669, GSM803401; PITX1: GSM1208667; NFE2: GSM1067276, GSM1427076, GSM935414, GSM935652; ASCL2: GSM1208591; GRHL2: GSM1125982, GSM1125983, GSM1125984, GSM1239569.We converted hg18 genomic positions to hg19 coordinates using the liftOver tool from UCSC (http://genome.ucsc.edu/cgi-bin/hgLiftOver). Where multiple data sets existed for a given factor, data set were merged to create a union peak set. Random peaks were generated maintaining the same chromosome proportion in the original peak file to create 100 iterations in each random peak simulation. To identify the number of genes associated to cistrome peaks we considered a distance window of 50 kbp from each TF peak coordinates using the distance() function from GenomicRanges package version 1.28.4. The P-value was calculated using the following expression: *P* = (r +1)/(n + 1), where r is the number of random values greater or equal than observation value and n is the number of random iterations.

### Data and code availability

All raw RNA sequencing data are also accessible at http://tcgabiolinks.fmrp.usp.br/OvarianRNAseq/. The code used to perform the analyses is available as an Rmarkdown at: http://tcgabiolinks.fmrp.usp.br/OvarianRNAseq/.

## Acknowledgements

The results shown here are in part based upon data generated by the TCGA Research Network: http://cancergenome.nih.gov/. Some of the normal tissue specimens were collected as part of the USC Jean Richardson Gynecologic Tissue and Fluid Repository, which is supported by a grant from the USC Department of Obstetrics & Gynecology and the NCT Cancer Center Shared Grant award P30 CA014089 (to the Norris Comprehensive Cancer Center). K.L. is supported in part by a K99/R00 Pathway to Independence Award from the NIH (R00CA184415), institutional support from the Samuel Oschin Comprehensive Cancer Institute at Cedars-Sinai Medical Center and a Career Development Award from The Tower Cancer Research Foundation. H.N. and M.A.S.F. are supported by grants 2015/07925-5 and 2017/08211-1 from Sao Paulo Research Foundation (FAPESP). H.N. is also supported by an institutional grant (Henry Ford Hospital). This work was supported in part by the Ovarian Cancer Research Fund Alliance Program Project Development Grant (373356): Co-Evolution of Epithelial Ovarian Cancer and Tumor Stroma. Additional support for this work came from NIH/NCI grants 1R01CA211707 and 1R01CA207456 and OCRF award 258807.

